# Hidden in the genomic bycatch: insights into genome reorganization and population structure of the parasitic nematode *Contortylenchus reversus*

**DOI:** 10.64898/2026.08.07.743562

**Authors:** Yahir De Jesus Campusano, German Lagunas-Robles, Lewis Stevens, Erik J. Ragsdale, Ryan R. Bracewell

## Abstract

Insect-parasitic nematodes are widespread and often significantly reduce host fitness, yet we know surprisingly little about most species. *Contortylenchus reversus* is a hemocoel-inhabiting parasitic nematode that infects *Dendroctonus* bark beetles, notably impacting host mobility and fecundity. We first detail a chromosome-scale genome assembly of *C. reversus*, recovered serendipitously from a sequencing project targeting a host (*Dendroctonus ponderosae*). We assembled the 79.4 Mb genome into nine linkage groups and, through transcriptome-aided annotation, identified 11,244 protein-coding genes. Synteny comparisons with the only relatives for which there are complete assemblies reveal extensive chromosomal rearrangements and extreme loss of gene collinearity suggesting these insect-parasitic nematodes may have exceptionally malleable genomes. Using this genome assembly and repurposed reduced-representation genomic data from 707 *D. ponderosae* individuals, we investigated infection frequencies and population structure, identifying infection rates ranging from 0% to 50% across 18 geographically widespread collection sites. Population structure of *C. reversus* appears broadly concordant with the structure of the host beetle, suggesting a shared evolutionary history, while genetic variation (nucleotide diversity) in the parasitic nematode is highly reduced in comparison to its host. These results offer insights into its population genetics, host associations, and the evolutionary dynamics of a nematode–beetle interaction and highlight how genomic bycatch can reveal previously hidden details about an important species in a complex community.

## INTRODUCTION

Genome sequencing projects of animals often unintentionally capture and sequence DNA from organisms other than the intended target (Vancaester and Blaxter 2023; Khalaf et al. 2025; Khalaf 2026). This is especially prevalent when working with small species, where DNA extraction is performed on whole individuals. As a result, a taxonomically diverse array of non-target DNA is often sequenced. However, only relatively small organisms with small genomes present in high abundance within a sample are likely to generate enough sequencing reads to allow for the assembly of their genomes. Such non-target assemblies typically include bacteria and other unicellular organisms, which are frequently treated as “bycatch” or contamination and are discarded; a practice that unfortunately results in many species being overlooked and additional genomic resources left undescribed. However, in theory other larger multicellular organisms could also be present as bycatch and assembled under certain sequencing scenarios.

During a high-coverage genome sequencing project targeting the Y chromosome of the mountain pine beetle, *Dendroctonus ponderosae* (Lagunas-Robles et al. 2026), a major forest pest that reproduces by killing pine trees, we recovered two complete genome assemblies from what we were later able to identify as the parasitic nematode *Contortylenchus reversus* (Thorne 1935) RÜHM 1956. The genus *Contortylenchus* belongs to the family Allantonematidae (Rhabditida: Tylenchomorpha), a broad group of nematodes known to parasitize numerous insect orders (Arthurs and Heinz 2003; Perlman et al. 2003; Poinar 2003; Poinar et al. 2004). *Contortylenchus reversus* was identified as the main hemocoel parasite of the mountain pine beetle (Thorne 1935) and of two other equally well-known pest *Dendroctonus* species, the Douglas-fir beetle (*D. pseudotsugae*) and the spruce beetle (*D. rufipennis*) (Massey 1974). Infection prevalence fluctuates annually and geographically, affecting up to 50% of beetles in some populations (Massey 1974). All life stages of the nematode, including gravid females, eggs, and the four larval stages, can be found coexisting within larval, pupal, or adult beetles. In some cases, a single beetle can host more than a thousand eggs and larvae (**Fig. S1**) (Thorne 1935; Furniss 1967; Thong and Webster 1973; Thong and Webster 1975b).

*Contortylenchus* species are oviparous with juvenile larvae being excreted by infected adult beetles into tunnels (galleries) that are excavated by the beetle while colonizing host trees (Massey 1974; Macguidwin et al. 1980a; Gibb and Fisher 1986). There, free-living juvenile males and females feed, develop and mate. During this free-living stage, *Contorylenchus* species likely enter a mycetophagous (fungi-feeding) phase where it joins a diverse community of bark beetle associates including other species of nematode, mites, and symbiotic (often mutualistic) fungi introduced into the tree by the beetle (Six 2012; Mercado et al. 2014; Hofstetter et al. 2015). Mating takes place in the beetle gallery, whereupon males then die, and fertilized females then penetrate the cuticle of developing beetle larva, usually in early instars. Inside the host, the nematode matures in synchrony with beetle development, eventually releasing offspring into the hemocoel and expelling them through the digestive tract into the gallery, completing the cycle. Typically, the nematode completes one generation per host generation (Massey 1974; Gibb and Fisher 1986). Infected adult beetles typically contain ∼10-20 adult female nematodes (Massey 1974).

Parasitic nematodes have been shown to significantly alter bark beetle physiology and behavior. For example, infections have been shown to impair beetle flight (Atkins 1961; Ashraf and Berryman 1970) and alter hemolymph composition and oocyte and testis development (Thong and Webster 1975a; Tomalak et al. 1990). More significantly and specifically, *C. reversus* infections in the Douglas-fir beetle have been found to reduce female fecundity by 33– 50% during tree colonization, resulting in shorter egg galleries (Massey 1974; Thong and Webster 1975b). The related nematode *C. brevicomi*, which infects the southern pine beetle (*D. frontalis*), can reduce fecundity up to 74% (Macguidwin et al. 1980b). Thus, parasitic nematodes, and *Contortylenchus* species specifically, clearly impact host beetle fitness and can influence beetle population dynamics (Massey 1974). Despite this important host–parasite interaction and potential for use in biological control, many insect-parasitic nematodes, including *C. reversus,* remain poorly studied and few genomic resources are currently available.

Here, we present a study using only “bycatch” genomics data to explore an important parasitic nematode. We present a chromosome-level genome assembly of *Contortylenchus reversus*, a representative of the Allantonematidae, a group of insect-parasitic nematodes that remains poorly understood across many ecological and evolutionary axes, and for which genomic resources are scarce. Through our broader comparative genomic analyses, which include several other species of Tylenchomorpha, we reveal that this group has a reduced and unique gene content that differs significantly from other nematodes. Furthermore, their genome structure is highly plastic and not typical of most nematodes. We also repurposed an existing genomic dataset targeting mountain pine beetles to investigate infection frequencies and nematode population structure. Our results provide valuable insight into infection prevalence and the spatial population structure of the nematode and beetle, offering a broader view of host– parasite dynamics and potential for tight coevolution in the system.

## RESULTS AND DISCUSSION

### Complete *Contortylenchus reversus* genome assembly and annotation

A combination of PacBio HiFi and Hi-C scaffolding allowed us to independently produce two nematode genome assemblies that were distinct from the host beetle and were broadly collinear (**Fig. 1A, Fig. S2**). Here, we detail the nematode assembly from Idaho male beetles, although results are qualitatively similar across the two assemblies as they were found to be similar in total length and lacked any sizable structural difference (**Fig. S2**). The Idaho assembly included nine linkage groups and was 79.4 Mb in total length (**Fig. 1A**, **Table 1**). An additional 11 contigs of putative nematode origin were identified but were unable to be placed and therefore left unassigned (**Table 1**). The average autosomal depth of long-read sequencing for the Idaho male beetle was 61×, while for the parasitic nematode(s) it was 48×, which may have contributed to the differences seen in the assembly contiguity (**Fig. 1A**). The comparable coverage of nematode and beetle genomes is likely due to the high parasite burden in these beetles, which can harbor hundreds or even thousands of nematode eggs and larvae, produced by several founder females. In the beetle (∼273.5 Mb genome), many chromosomes were assembled in a single contig (Lagunas-Robles et al. 2026), while for the nematode the average number of contigs per linkage group was seven (**Table 1**). The canonical nematode telomere sequence (TTAGGC)_n_ was identified on both ends of five linkage groups (LG3, LG4, LG6, LG7, LG8) and on one end of the other four. One single-end LG from the Idaho assembly (LG1) was found to have telomere on both ends in the second assembly (Nevada), and therefore a total of six of nine linkage groups were found to be complete with telomeres on both ends. Hi-C data did not indicate that the three single-end LGs (LG2, LG5, LG9) are any part of a single chromosome as there was no enrichment of contacts between those linkage groups (**Fig. 1A, Fig. S2**). Thus, these assemblies are of high quality and are chromosome-level.

**Figure 1.**
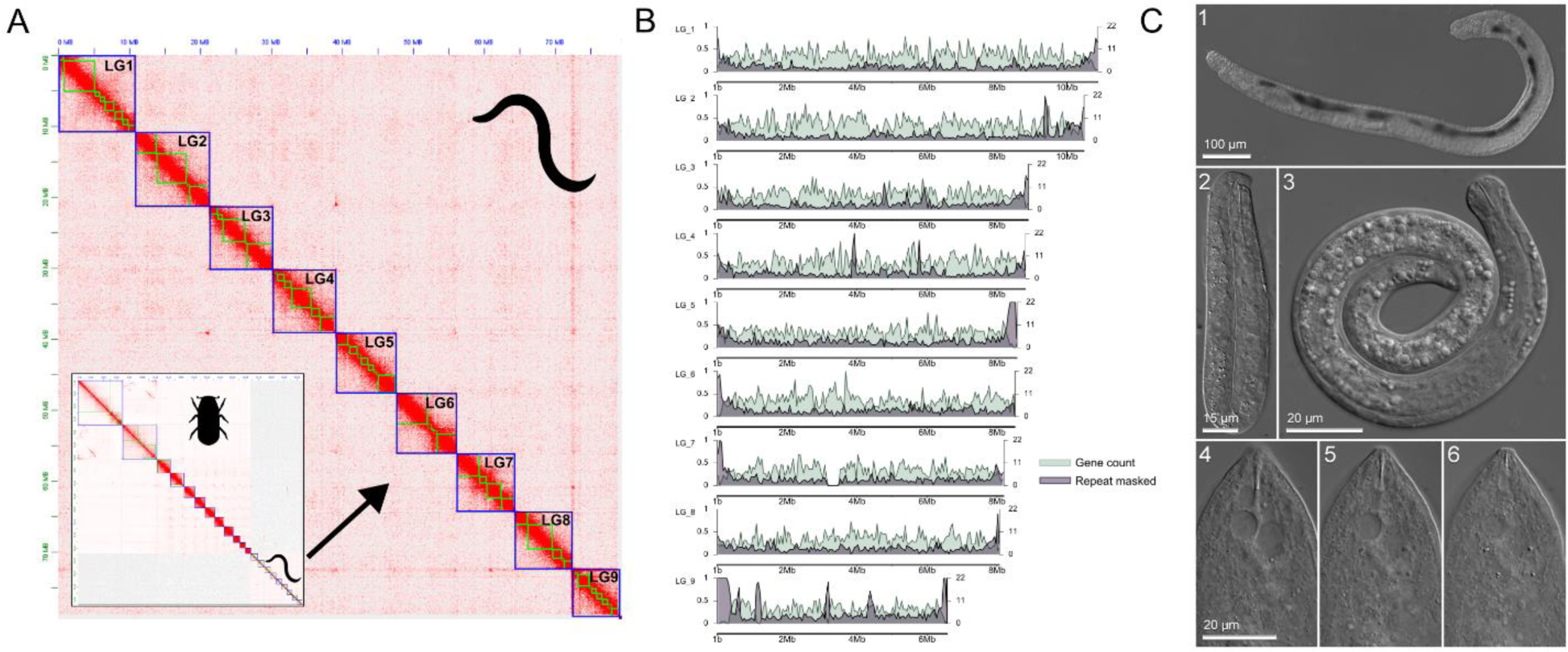
Chromosome-level genome assembly of *Contortylenchus reversus* recovered as bycatch during sequencing of host beetle. **A**) Hi-C contact map showing nine linkage groups of the beetle parasite, *C. reversus*. Blue boxes outline scaffolded linkage groups, green boxes identify contigs. Inset shows combined Hi-C contact map of host beetle *Dendroctonus ponderosae* and *C. reversus*. **B**) Gene counts (right axis) and proportion of masked repeats (left axis) in 50 kb windows across the *C. reversus* genome assembly. **C**) Morphological identification and micrographs of *C. reversus* isolated from *D. ponderosae*: (1) parasitic, egg-laying female, (2) ensheathed second-stage larva, (3) third-stage larva, (4-6) head of parasitic female in three focal planes, showing the stylet (4) knobs, (5) shaft, and (6) cone. Photos: Erik Ragsdale.

**Table 1.** Genomic features of *Contortylenchus reversus.* General genome assembly and annotation characteristics including the number of scaffolding locations (stitch points), and results from repeat masking.

| Linkage group | Length (bp) | Stitch points | GC content (%) | Protein coding genes | Masked nucleotides (bp) | Masked (%) |
| --- | --- | --- | --- | --- | --- | --- |
| LG_1 | 10,888,573 | 8 | 35.96 | 1,643 | 1,437,863 | 13.21 |
| LG_2 | 10,488,831 | 6 | 35.99 | 1,560 | 1,397,101 | 13.32 |
| LG_3 | 8,894,349 | 4 | 35.93 | 1,266 | 1,243,321 | 13.98 |
| LG_4 | 8,816,446 | 5 | 36.01 | 1,331 | 1,235,216 | 14.01 |
| LG_5 | 8,570,856 | 9 | 35.61 | 1,157 | 1,432,719 | 16.72 |
| LG_6 | 8,513,501 | 5 | 35.97 | 1,278 | 1,284,262 | 15.09 |
| LG_7 | 8,196,639 | 3 | 35.31 | 1,096 | 1,224,352 | 14.94 |
| LG_8 | 8,071,758 | 4 | 35.74 | 1,093 | 1,178,483 | 14.60 |
| LG_9 | 6,572,245 | 12 | 36.75 | 818 | 1,719,699 | 26.17 |
| Unassigned | 385,502 | 0 | 43.37 | 2 | 382,859 | 99.31 |

Our initial assessments of genome assembly completeness revealed that out of 3,131 total nematode BUSCOs, only 2,271 (72.5%) genes were complete, while 760 (24.3%) were entirely missing. The GC content was found to be rather uniform across chromosomes, fluctuating around 35–36%; a range typical of many nematodes. Our identification of repetitive elements was also found to be rather typical for nematodes given the size of the genome assembly, with 15.9% of total bases being identified as being derived from repetitive sequence with the major components being unclassified repeats (∼8%), DNA transposons (∼4%), and retroelements (∼1%). The proportion of masked bases was relatively uniform across most chromosomes, with values between 13-17%. However, LG_9 stood out as atypical with a significantly higher percentage of masked bases (26%) (**Table 1**). A transcriptome-guided annotation, generated from bycatch RNA-seq data originally targeting the host beetle, identified 11,244 protein-coding genes which appear broadly distributed across all chromosomes (**Fig 1B**). LG_9, which was identified as the most repetitive, also was found to contain the fewest genes (**Table 1**). The somewhat uniform gene and repeat content found across all *C. reversus* chromosomes is occasionally seen in some nematode species (Woodruff and Teterina 2020), although the more common pattern in many nematodes is for chromosome centers to be enriched for genes and depleted of repeats, while the inverse pattern occurs on the ends of the chromosomes.

Because the nematode genomes were originally sequenced without morphological vouchers, we dissected new specimens of *D. ponderosae* in an attempt to reisolate the same species. Some beetles collected from an Idaho lab colony were positive for *Contortylenchus*, whose characters agreed with published descriptions of *C. reversus* (**Fig. 1C**). We then sequenced a fragment of the 18S rRNA gene from individual adult parasitic females, which had an exact match to the homologous sequence fragment extracted from the assembled *C. reversus* genome, definitively linking vouchers of this morphospecies to the genome assembly.

### Nematodes of Tylenchomorpha have unique gene content and malleable genomes

The low completeness score and missing BUSCOs in *C. reversus* led us to explore if this was restricted to this species and/or potentially a genome assembly issue, or was a broader feature of this group of nematodes. We therefore analyzed published genomes of several species of Tylenchomorpha: three with high-quality assemblies, at or approaching chromosome-level, including the soybean-cyst nematode, *Heterodera glycines* (Masonbrink et al. 2021), a carrot weevil nematode, *Bradynema listronoti* (Ste-Croix et al. 2022), and a bumblebee nematode *Sphaerularia bombi* (GenBank accession: GCA_964235305.1). We also explored five additional tylenchids of variable assembly contiguity, including *Deladenus siricidicola* (GCA_009724625.1), *Ditylenchus destructor* (GCA_043789845.1), *Ditylenchus dipsaci* (GCA_004194705.1) *Subanguina moxae* (GCA_000981365.1) and *Anguina tritici* (GCA_029339935.1); the model *C. elegans* was also included to help establish evolutionary relationships and explore gene content. We first constructed a phylogeny of these species using 1,046 shared BUSCOs. This analysis revealed well-supported relationships that challenged more weakly supported groups from previous analyses based only on rRNA genes (van Megen et al. 2009; Takahashi et al. 2024), although direct comparison is hindered by differences in different taxon representation. We then used our inferred tree for comparative analyses (**Fig. 2A**).

**Figure 2.**
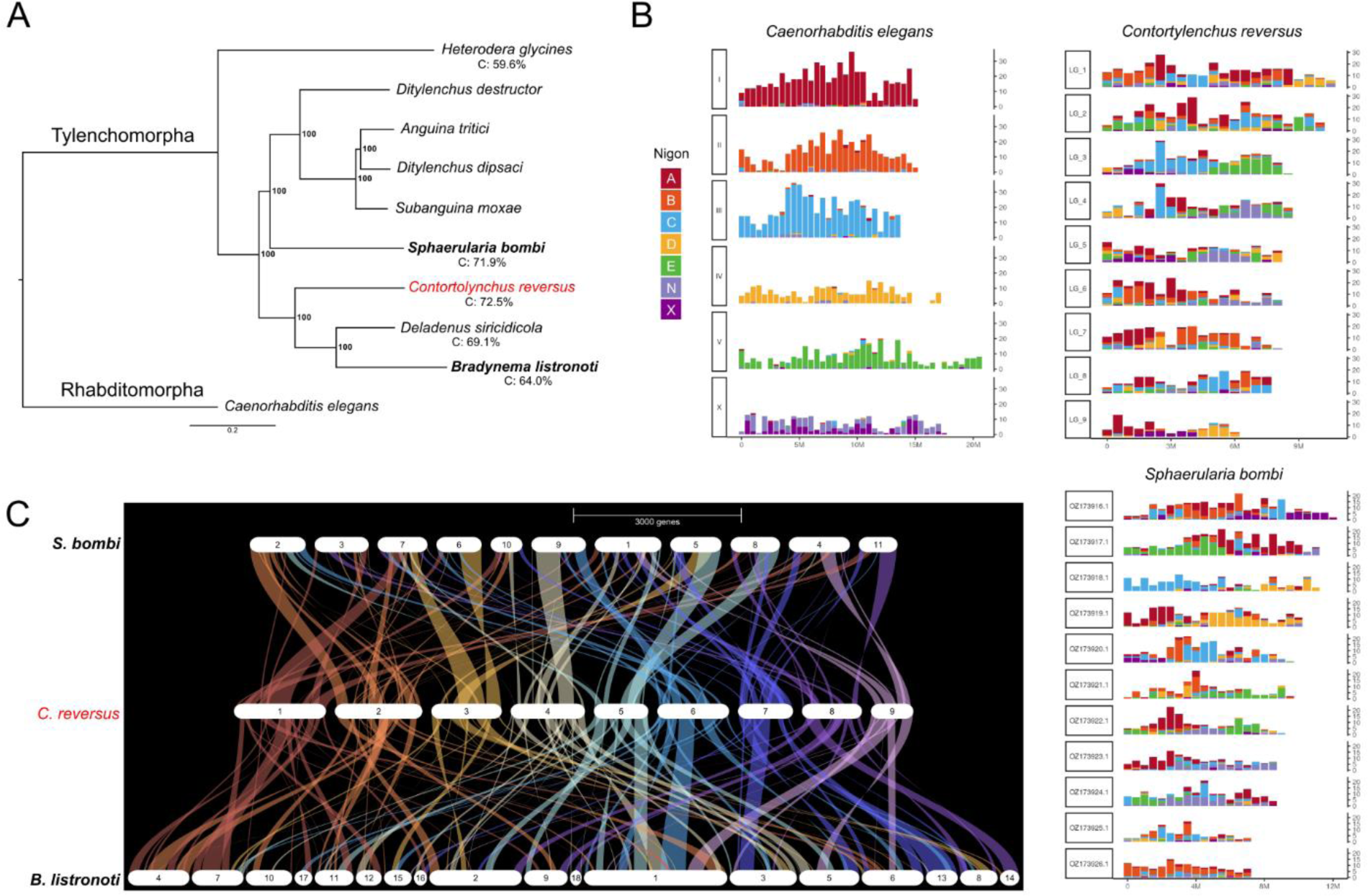
Tylenchid genomes are missing numerous nematode BUCSOs and are structurally malleable. **A**) Phylogenetic relationships among nematodes with high-quality genome assemblies. BUSCO genome completeness percentage for select species with high-quality and well characterized genomes shown below. **B**) Chromosomal locations of Nigon element-associated BUSCOs in three genomes; the model *C. elegans* and insect-parasitic *C. reversus* and *S. bombi*. In *C. elegans*, each chromosome largely corresponds to a single Nigon element (N and X fused in ancestor). **C)** GENESPACE riparian plot showing lack of chromosome conservation between insect-parasitic nematodes *S. bombi*, *B. listronoti*, and *C. reversus* (center, reference). Note that the *B. listronoti* assembly is not chromosome-level (Ste-Croix et al. 2022).

We found that, across the four species with the highest-quality genome assemblies, all had low completeness scores, with *S. bombi* comparably low (71.9%), while *B. listronoti* (64.0%) and *D. siricidicola* (69.1%) displayed slightly lower yet similar values to *C. reversus* (**Table S1**) *Heterodera glycines* reported the lowest proportion of complete genes (59.6%) (**Table S1**). Thus, this reduced gene set appears to be a feature of Tylenchomorpha (**Fig. 2**) and many of the genes from the BUSCO orthodb10 database appear to be broadly missing from Tylenchomorpha genomes, as pointed out by others (Masonbrink et al. 2021). We found the *C. reversus* genome size was similar, but slightly smaller, than other related nematode species (*H. glycines* = 157.98 Mb; *S. bombi* = 109.1 Mb*, B. listronoti* = 80.64 Mb, **Table S2**). Interestingly, *C. reversus* had the fewest genes at 11,244, and *S. bombi* was similarly low (11,673), while *B. listroni* (17,886), *H. glycines* (22,465), and *C. elegans* (19,984) have substantially more genes (**Table S2**). However, comparisons of gene number across species are challenging given different annotation pipelines, stringency, and curation. In total, we found C. *reversus* to have a small genome and reduced number of protein coding genes, but genomically does not appear entirely outside the norm for this group of nematodes.

To characterize chromosome structure and genome stability through time, we next explored genome synteny (chromosome conservation or lack of *inter*chromosomal rearrangement) and collinearity (gene order or *intra*chromosomal rearrangement) with GENESPACE (Lovell et al. 2022) and the four species with chromosome (or near chromosome) level assemblies. The last common ancestor of rhabditid nematodes had seven chromosomes, which have been termed Nigon elements (Tandonnet et al. 2019; Gonzalez de la Rosa et al. 2020). In many species, such as *C. elegans*, these elements are largely conserved while in other species occasional chromosomal fusions/fissions have been described leading to rearrangement of this ancestral configuration (Gonzalez de la Rosa et al. 2020; Kounosu et al. 2024). Surprisingly, our results showed a remarkable loss of both synteny and collinearity between these species and a complete breakdown of Nigon elements (**Fig. 2B-C**., **Fig. S3**). For example, LG2 in *C. reversus* aligns partially with chromosome 2 in *S. bombi* but also shares blocks with chromosomes 7 and 10. Similar patterns are observed for all other chromosomes in *C. reversus*, which show split synteny across multiple chromosomes in *S. bombi* (**Fig. 2C**). *Bradynema listronoti* exhibits an even more fragmented synteny pattern with *C. reversus* (**Fig. 2C**). To further explore this extreme breakdown of synteny, we tracked the physical locations of 1:1 orthologs shared between species and found not only changes in synteny, but also extensive reshuffling of gene order and loss of collinearity (**Fig. S4**). In comparison with *B. listronoti*, orthologs appeared widely scattered across multiple linkage groups in *C. reversus*, showing no clear one-to-one chromosomal correspondence. A similar pattern was observed in comparison with *S. bombi*, where orthologs were distributed across multiple chromosomes in both species and were highly shuffled (**Fig. S4**). These patterns of genome evolution suggest numerous ancestral chromosomal fusions likely occurred followed by the proliferation of chromosomal inversions and intrachromosomal gene movement. Although what we document is uncommon in nematode genomes, a rearrangement of Nigon elements driven by chromosomal fusions has been observed in other nematode lineages, such as in *Strongyloides* (Kounosu et al. 2024) and *Diploscapter* (Chung et al. 2026). However, a unique feature of *C. reversus* and *S. bombi* appears to be subsequent unique fissions that fragmented these ancestrally fused/shuffled chromosomes; a series of evolutionary events that have dramatically reshaped genome structure and gene order across this group of nematodes.

### Contortylenchus reversus is geographically widespread and has correlated population genetic structure with its host beetle

Beetles infected with *C. reversus* appear to carry a large load of both adult and juvenile individuals (Massey 1974)(**Fig. S1**). We reasoned that previous population genomic analyses aimed at the beetle *D. ponderosae* may have contained infected individuals, and thus sufficient nematode genomic reads, to 1) estimate infection frequencies from geographically widespread populations and 2) and explore nematode population structure. Although numerous studies have examined population genetic structure of various *Dendroctonus* beetles (Maroja et al. 2007; Anducho-Reyes et al. 2008; Ruiz et al. 2010; Dowle et al. 2017; Bracewell et al. 2018; Havill et al. 2019) and some studies have characterized correlated population structure of mutualistic symbiotic fungi (Roe et al. 2011; Bracewell et al. 2018), our understanding of nematode genetic structure in these complex communities has yet to be explored.

We reanalyzed data originally produced to explore population structure in *D. ponderosae*, a reduced representation genomic dataset (ddRAD-seq) from 707 beetle DNA extractions from 35 locations across the western US, southern Canada, and northern Baja California, Mexico (Dowle et al. 2017). We first mapped all sequencing reads to our repeat-masked *C. reversus* genome assembly and using a K-means clustering approach of total read counts, separated nematode-infected from uninfected samples (**Fig. S5**). Rangewide, we found the infection rate varied across the 18 locations where nematodes were detected (**Fig. 3A**, **Table S3**). The highest prevalence was found in Arizona (AZ1), where 50% of beetles appeared to harbor nematodes. Other heavily infected locals were found in several geographically distant locations, including Idaho (ID4, 35%), Utah (UT3, 31.6%), and California (CA3-4, 32.3%). Moderate levels were observed from a collection in Nevada (NV5, 25%), while other regions CO, NV2, and WY, showed relatively low infection rates (≤ 5.3%) (**Fig. 3A**, **Table S3**). At 17 locations, we failed to confidently detect *C. reversus*, although there was no clear geographic pattern or obvious ecological pattern (e.g., host tree species), connecting uninfected populations (Dowle et al. 2017). Interestingly, these frequencies appear consistent with previous geographically restricted studies exploring infection frequencies in *D. ponderosae* and its relative, *D. pseudotsugae* (Massey 1974). However, it is important to note that various factors could influence our measures including the beetle life stage, tissue composition/bias in DNA extraction, and our stringent read count-infection status cutoff. Therefore, our results are likely a conservative estimate of infection frequencies, but nonetheless, demonstrate *C. reversus* is a consistent and widespread parasitic partner of *D. ponderosae* with significant impacts on its ecology.

**Figure 3.**
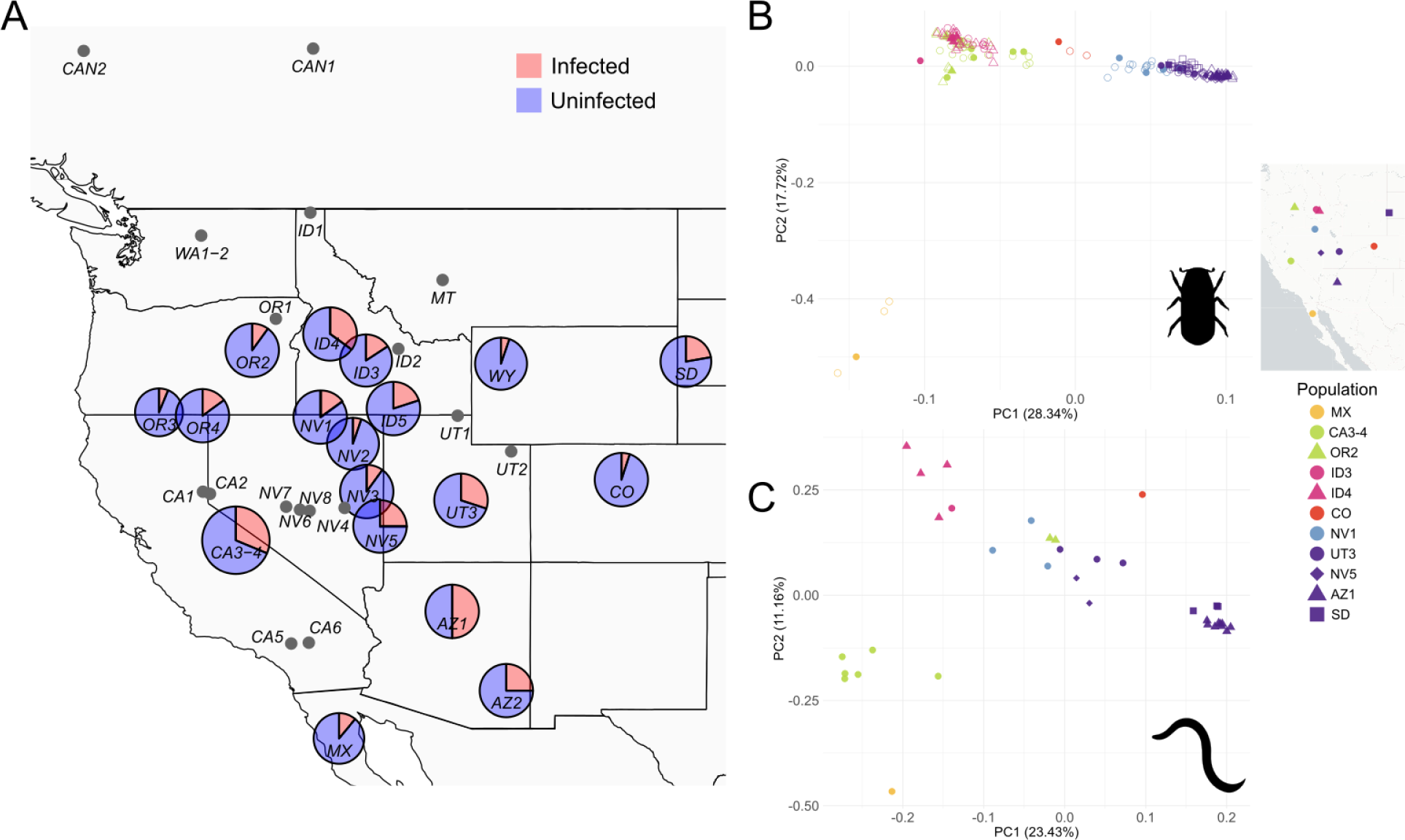
Repurposed genomics data shows *Contortylenchus reversus* is geographically widespread, varies in local abundance, and has correlated population genetic structure with host beetle *Dendroctonus ponderosae*. **A)** Collection locations of host beetle from western North America with inferred proportions of beetles infected with *C. reversus*. For locations where beetles were found to be infected with nematodes, pie charts are scaled by the total number of individuals tested (**Table S3**). Locations represented by grey points had zero infections. Note ID3 and ID4 were slightly shifted for visualization. **B**) Principal components analysis (PCA) of 1,467 autosomal SNPs from the host beetle *D. ponderosae*, with points colored by previously identified population genetic clusters (Dowle et al. 2017). Solid symbols identify nematode-infected beetles while open symbols denote other individuals from the same collection location. C) PCA of 187 SNPs of *C. reversus*, with samples color-coded as in **B**).

Given the broad geographic association, and intricately linked life history, we next explored if the population structure of the nematode matched that of host *D. ponderosae*. We first identified 35 nematode samples with sufficient sequencing coverage for SNP identification and then explored population genetic structure of those specific beetle populations using both infected and uninfected high-quality individuals (163 total). We followed an exceptionally stringent SNP filtering approach (see METHODS) to ensure only beetle derived SNPs were analyzed, resulting in the identification of 1,467 autosomal coding sequence SNPs found across 1,316 *D. ponderosae* genes representing all 11 autosomes. Our principal component analysis (PCA) of beetle SNP data showed population genetic structure consistent with previous studies (Mock et al. 2007; Bracewell et al. 2017; Dowle et al. 2017). The first two principal components explained 28.34% (PC1) and 17.72% (PC2) of the total genetic variation. Samples from Arizona, Nevada, Utah and South Dakota (SD, AZ1, NV5, NV1, UT3) formed a distinct cluster, a grouping consistent with a cryptic species identified via Y chromosome type (Bracewell et al. 2017; Dowle et al. 2017). Colorado, Idaho, Oregon and California samples clustered by population, but had less clear geographic structure, a result of ongoing autosomal gene flow via introgression between two other cryptic beetle species (Bracewell et al. 2017; Dowle et al. 2017). Samples from Mexico (MX) stand out as very distinct, likely due to being from a small and isolated population at the southernmost range limit of *D. ponderosae*. Nematode infected beetles were found to cluster with uninfected individuals from the same collection location (**Fig. 3B**).

We next explored the population structure of the nematode using 187 high-quality SNPs with 101 SNPs located in the coding sequence of 101 genes found across all nine linkage groups. We examined the remaining 86 SNPs by mapping a 400 bp region containing each SNP against the *D. ponderosae* genome. None of the nematode SNP regions mapped to the beetle genome, confirming these SNPs were indeed of nematode origin. It is important to note that SNPs were not identified from a single nematode, but likely a mix of mostly related juveniles contained within the host beetle. We found a striking similarity in our PCA where nematode structure broadly reflected host beetle structure (**Fig. 3C**). The first two principal components explained 23.43% (PC1) and 11.16% (PC2) of the total genetic variation (**Fig. 3C**). Notably, samples from Arizona and South Dakota (AZ1, SD), two geographically distant locations with genetic similarity in beetles (**Fig. 3B**)(Dowle et al. 2017), show clear clustering in the nematode. Similarly, the Mexico sample again stood out as distinct although closest to California samples (CA3-4). A Mantel test based on the Euclidean distances between PC1 and PC2 scores of paired samples, both beetle and nematode present in their respective PCAs (*n* = 34, permutations = 1,000), revealed a highly significant correlation (r = 0.60, p < 0.001). Overall, nematodes showed clear clustering by population and patterns consistent with isolation-by-distance and known phylogeographic patterns described in the host beetle; patterns that are largely the result of historical host tree distributions and Pleistocene glacial refugia, cryptic species boundaries, and current forest connectivity (Mock et al. 2007; Bracewell et al. 2017; Dowle et al. 2017). Interestingly, despite this shared evolutionary history, the nematode harbors nearly 17-fold lower levels of genetic variation (**Table S4**, nematode nucleotide diversity (π) = 0.03% ± 0.007) compared to the host beetle (**Table S5**, autosomal π = 0.5% ± 0.09), revealing substantial differences in their respective effective population sizes.

The striking pattern of correlated population genetic structure between host beetle (*D. ponderosae*) and parasitic nematode (*C. reversus*) suggests a tightly shared evolutionary history between these two species. This finding is somewhat surprising given that two other *Dendroctonus* species are also considered hosts for the morphospecies *C. reversus*, yet they have different geographic ranges and reproduce in host tree species well outside the host-breadth of the *Pinus* generalist *D. ponderosae* (Six and Bracewell 2015). As the names suggest, the Douglas-fir beetle reproduces primarily in Douglas-fir (*Pseudotsuga menziesii*) and the spruce beetle in several species of spruce (*Picea glauca*, *Picea engelmannii*, *Picea sitchensis*, *Picea rubens*). Pines, spruces, and Douglas-fir do co-occur in mixed conifer stands throughout parts of western North America, but these three species of beetle rarely attack, and are incapable of successfully reproducing in non-host trees. Thus, the host tree specificity exhibited by different *Dendroctonus* beetle species should thereby limit the possibility of horizontal movement of parasitic nematodes between species, even during any free-living life stage. It is therefore plausible that *Contortylenchus reversus* is actually a complex of cryptic species, given the difficulty of delimiting closely related nematode species using morphological characters alone. Each cryptic species is likely tightly coevolving with its host beetle species during the parasitic stage, and more loosely adapting/coevolving during the free-living stage with a unique community of mites, commensal nematodes, and symbiotic fungi associated with each *Dendroctonus* beetle species (Mercado et al. 2014; Hofstetter et al. 2015). These putative cryptic *Contortylenchus* species await further study to establish species boundaries and to fully understand their impacts in these complex communities which collectively shape western North American forests.

## CONCLUSIONS

We demonstrate that high-coverage sequencing of a target species, combined with careful analysis of bycatch data, can yield genome assemblies for additional eukaryotic taxa. Further, by repurposing existing genomic datasets we can generate valuable insights into some of the planet’s lesser-studied organisms. Insect-parasitic nematodes are diverse and widespread, yet their basic biology, ecology, genome structure, and evolutionary history remain poorly understood for most species. Further, the approach presented here opens new avenues for studying these important members of complex ecological communities, as exemplified by the tree–bark beetle–symbiont systems that have far-reaching ecological and economic impacts. Leveraging such datasets enables population genomic analyses of organisms that have traditionally been overlooked, while also facilitating the investigation of shared genetic structure across interacting species. These opportunities may ultimately provide a deeper understanding of how interspecific interactions shape patterns of genetic variation and population structure across communities.

## METHODS

### Raw genomic data

For a detailed description of the original beetle collection locations and PacBio HiFi, Hi-C and RNA-seq data generation see (Lagunas-Robles et al. 2026). What we report here is a brief summary specific to the nematode *C. reversus*. The two collection locations and associated datasets in (Lagunas-Robles et al. 2026) are from the Idaho male *Dendroctonus ponderosae* and Nevada female assembly (JBTYYW000000000 and JBTYYX000000000, respectively). The PacBio HiFi, Hi-C, and RNA-seq data were generated from different beetles at each location.

### Genome assembly and Hi-C scaffolding

Raw PacBio Hifi reads were assembled using Hifiasm v0.19.8-r603 with the option –primary (Cheng et al. 2021). To verify the assembly and evaluate base-level accuracy, we re-mapped our raw PacBio Hifi reads to the primary assembly using minimap2 v2.28 with option -ax map-pb (Li 2018; Li 2021). All downstream sorting and indexing of mapped reads were done using samtools v.1.20 (Danecek et al. 2021). To scaffold contigs in our assembly, raw paired-end Hi-C reads were mapped using BWA mem v0.7.17 (Li and Durbin 2009). We used Picard v 3.1.1 to process bam files, and AddOrReplaceReadGroups to standardize group information and MarkDuplicates to identify duplicate reads. After removing the duplicates, we used Bedtools v2.31.1 (Quinlan 2014) to create a BED file from the Hi-C read alignments based on the mapped BAM file Hi-C contact intervals, which was required as input for YaHS 1.2.2 (Zhou et al. 2023). YaHS 1.2.2 was used to produce the Hi-C contact matrix and scaffold the genome assembly. To visualize and manually edit the scaffolded assembly, we used Juicebox v2.20 (Durand et al. 2016a). Pre-processing was done using JuicerTools v3.0 (Durand et al. 2016b).

### Estimating genome coverage and identifying unplaced nematode contigs

Per-base coverage for all scaffolds and contigs in the beetle + nematode assembly was determined using bedtools with genomecov and the -g option (Quinlan 2014). Genomic coverage was calculated in 50 kb windows and then plotted using Rstudio v. 2024.12.0+467 and dplyr library version 1.1.4. To identify and retain any putative nematode contigs not placed via Hi-C scaffolding, yet avoid other contaminant contigs (e.g., bacteria, yeast) we first calculated the per contig coverage and flagged putatively nematode-derived contigs if the contig was ± 10× the scaffolded nematode assembly median. We then BLAST this subset of contigs against the entire nucleotide database and removed any with top hits to beetles, fungi, or bacteria.

### Genome annotation

Repetitive elements were first identified using RepeatModeler v2.0.5 (Flynn et al. 2020) and RepeatMasker v4.1.5 (Smit et al. 2013-2015) was used to create a softmasked assembly for annotation. To identify protein coding genes, we used transcriptomic data (SAMN61658811) from a nematode-infected beetle to generate input transcripts for the Braker pipeline v3.0.3 (Gabriel et al. 2024). Genome completeness was determined using BUSCO v5.7.1 (Simão et al. 2015) and the nematoda_odb10 database. For BUSCO comparisons with other relatives, we analyzed *Bradinema listronoti* (GCA_024678965.1), *Sphaerularia bombi* (GCA_964235305.1), *Deladenus siricidicola* (GCA_009724625.1), *Heterodera glycines* (GCA_004148225.2), and the model organism *Caenorhabditis elegans* (GCA_000002985.3).

### Genome characterization

To explore the genomic distribution of protein-coding genes and repetitive sequences in the *C. reversus* genome, we created a hard-masked genome assembly (above) and calculated the proportion of bases masked in non-overlapping 50 kb windows. Gene distribution was assessed by counting the number of protein-coding genes that fell in non-overlapping 50 kb windows. Gene and repeat abundance across each linkage group were visualized in KaryoplotR (Gel and Serra 2017).

### Nematode identification

For dissection, adult beetles were first rinsed in M9 buffer to exclude nematodes from the outside of the body, after which they were dissected in M9 buffer. If positive for tylenchid parasites, the hemocoel of the beetle contained ∼4-8 mature, adult parasitic females and hundreds of larvae of multiple life stages. Adult parasitic females were identified morphologically as belonging to the genus *Contortylenchus* based on its dorsally arcuate body and association with the hemocoel of bark beetles (Rühm 1956; Siddiqi 2000). Species identification followed (Thorne 1935) and (Thong and Webster 1973). Specimens agreed with published descriptions of *Contortylenchus reversus* in possessing a stout parasitic female, small relative to other *Contortylenchus* species, with a strongly dorsally recurved anterior body, a reflexed ovary exceeding the body length, and a conoid tail terminating in a distinct mucro (**Fig. 1C**), as well as being recovered from *D. ponderosae*, consistent with the nematode’s original description. For molecular barcoding of morphological *C. reversus* vouchers, primers were designed for the 18S gene based on the assembled genome (forward primer, GAAGATTAAGCCATGCATGC; reverse primer, AAACTTGGCAATTGCTTTCG).

### Synteny comparisons

To explore genome synteny and collinearity among *C. reversus* and related nematodes we used GENESPACE (Lovell et al. 2022). Species were selected based on assembly quality and completeness and/or putative close evolutionary relationship to *C. reversus*. Genome assemblies and annotations were retrieved from WormBase ParaSite (WBPS19) (Howe et al. 2017) and GenBank. We used *C. elegans* (GCA_000002985.3), *B. listronoti* (GCA_024678965.1, (Ste-Croix et al. 2022)), *H. glycines* (GCA_004148225.2, (Masonbrink et al. 2021), and *S. bombi* (GCA_964235305.1). For *S. bombi*, there was no available annotation, so we used Braker3 (Gabriel et al. 2024) along with the predicted proteins from *C. reversus* to annotate the genome. We followed the GENESPACE v1.2.3 pipeline, which uses DIAMOND v2.0.15 (Buchfink et al. 2021) for protein sequence alignment and performs BLASTP searches (Altschul et al. 1990; Camacho et al. 2009) to identify homologous proteins with an e-value threshold of 0.001. GENESPACE also utilizes OrthoFinder v2.5.4 (Emms and Kelly 2019) to cluster genes into orthogroups, identifying orthologous relationships. We used default settings to produce GENESPACE riparian plots. In addition, we visualized all 1:1 orthologues between *C. reversus* and each species listed above using RIdeogram (Hao et al. 2020).

### Phylogenetic tree

To explore evolutionary relationships, we used our previously identified BUSCOs (above) to construct a phylogeny using a supermatrix approach. We extracted the protein sequences using busco2fasta.py (available at https://github.com/lstevens17/busco2fasta) with the flags -- supermatrix_only and --psc 90. We identified 1,046 BUSCOs that were single copy and present in at least 90% of the genomes. These protein sequences were aligned using MUSCLE (v5.1) (Edgar 2004) and trimmed the alignments using trimAl (v1.4.rev15) (Capella-Gutiérrez et al. 2009) with the automated1 default option, which automatically selects optimal trimming parameters based on the input data. The final concatenated supermatrix contained 491,901 amino acids. We used this alignment to infer a species phylogeny using IQ-TREE (v.2.4.0) (Minh et al. 2020) under automatic model selection (-m MFP) with 3,000 ultrafast bootstrap replicates (-bb 3000) and 3,000 SH-aLRT tests (-alrt 3000). In addition to *C. reversus*, *S. bombi*, *B. listronoti*, *C. elegans*, and *H. glycines*, the phylogeny included the insect-parasitic *Deladenus siricidicola* (GCA_009724625.1), as well as the plant-parasitic *Ditylenchus destructor* (GCA_043789845.1), *Ditylenchus dipsaci* (GCA_004194705.1), *Subanguina moxae* (GCA_000981365.1), and *Anguina tritici* (GCA_029339935.1).

### Estimating infection frequency in beetle populations

We used published RAD-seq data generated from DNA extracted from the thorax of 707 *Dendroctonus ponderosae* individuals (Dowle et al. 2017). Raw sequencing reads were demultiplexed and filtered using process_radtags from the Stacks pipeline (Catchen et al. 2013), specifying the EcoRI restriction enzyme, Phred + 33 encoding, and the -q and -c flags to discard reads with low quality (Phred < 20) or any uncalled bases. The demultiplexed reads were mapped to the repeat-masked *C. reversus* genome using bwa v0.7.17-r1198-dirty (Li and Durbin 2009) and were sorted using samtools v1.17. Reads with a mapping quality ≥20 were counted using samtools view (Li et al. 2009).

To identify infected individuals based on mapped read counts, counts were first log2- transformed, and K-means clustering (k = 2) was applied to explore separation of uninfected and infected individuals. Based on a computed decision boundary a conservative coverage threshold of 4,096 mapped and filtered reads (Log2 (mapped reads) ≥ 12) was used to classify infection status. Visualization of infection frequencies among populations was performed in RStudio (v2024.12.0+467) using the packages ggplot2 (Wickham 2016), ggmap (Kahle and Wickham 2013), and scatterpie (Yu and Yu 2018).

### Nematode and beetle population genetic structure

We first used 35 individuals with the highest nematode read counts (above) to identify 11 focal nematode/beetle populations. We aligned the MPB data from 225 *Dendroctonus ponderosae* individuals from the focal populations to the ID male reference genome (Lagunas-Robles et al. 2026), JBTYYW000000000) with bwa v0.7.17-r1198-dirty (Li and Durbin 2009) and bams were sorted using samtools v1.17 (Li et al. 2009). We called all sites with reads ≥20 mapping quality in bcftools v1.17 (Danecek et al. 2021). We tried several filters in VCFtools v.0.1.16 (Danecek et al. 2011) and settled on the following to maximize high-quality SNPs from as many samples as possible. We generated a VCF file that only contained autosomal bi-allelic SNPs (--min-alleles 2, --max-alleles 2), removed indels (--remove-indels), and retained SNPs that were 100 bp up/downstream from EcoRI cut sites (--bed), but excluded SNPs that were in, or 5bp from, EcoRI recognition motif (--exclude-bed). We then retained SNPs that were present in ≥50% of samples (--max-missing 0.50), where the alternative allele was supported by at least three copies (--mac 3), had a minimum depth of 5 (--minDP 5), and thinned SNPs so each SNP was ≥ 1000 bp apart (--thin 1000). We retained SNPs in coding sequencing (CDS) regions using the ID male genome annotation (Lagunas-Robles et al. 2026). We identified 1,467 high-quality beetle CDS SNPs found across 1,316 genes that represented all 11 autosomes.

We called all sites and filtered nematode SNPs using the same pipeline described above, except we retained SNPs where the alternative allele was supported by at least two copies (--mac 2). This resulted in 187 high-quality nematode SNPs. To further confirm that the SNPs were indeed of nematode origin, we first determined how many resided in nematode CDS based on our annotation. Over half of the SNPs were present in worm CDS (101/187 high-quality SNPs). The remaining 86 SNPs were examined by extracting the neighboring flanks 199 bp regions flanking each SNP (400 bp total) using bedtools v2.31.0 getfasta (Quinlan 2014) and then using blastn v2.17.0+ (Camacho et al. 2009) to search against the ID male beetle genome (JBTYYW000000000). None of the 86 SNP regions mapped to the beetle genome.

To assess the population structure within the beetle and the nematode, we conducted a principal components analysis in plink v1.9.0-b.8 (Purcell et al. 2007) and specified a maximum missing genotype rate of 60% (--mind 60). The PCAs included 163 high-quality beetles and 35 high-quality nematode samples. Results were visualized in ggplot2 (Wickham 2016). We transformed the beetle and nematode PC1 and PC2 into euclidean distance matrices. To maintain pairwise relationships, we used 34 nematodes as one beetle lacked sufficient coverage to be included in the beetle PCA. Correlations were determined with a mantel test in the R package vegan v2.7-5, computed using Pearson’s product-moment correlation and tested for statistical significance using 1000 permutations.

We estimated nucleotide diversity (π) with pixy v2.0.0.beta14 (Korunes and Samuk 2021) for autosomes and linkage groups in the high-quality beetles (*n* = 163) and nematodes (n = 35), respectively, using an all-sites VCF. We generated an all-sites VCF with SNPs and merged invariant sites for RADtag loci. We filtered for autosomal bi-allelic SNPs, removed indels, and were 100 bp up/downstream from EcoRI cut sites, but excluded EcoRI recognition motifs plus 5 bp of adjacent upstream and downstream sequence. Retained SNPs were present in ≥50% of samples (--max-missing 0.50), with at least three copies supporting the alternative allele (--mac 3), had a minimum depth of 5 at each SNP (--minDP 5) and maximum depth of 500 per sample (--max-meanDP 500). Invariant sites (--max-maf 0) were filtered in the same manner prior to merging. We found these methods produced beetle π estimates that were largely consistent with previous whole genome sequencing estimates (Bracewell et al. 2017).

## ACKNOWLEDGEMENTS

We thank Dwayne Tally for assistance with several comparative genomic analyses. We also thank Jim Vanydgriff and Matt Hansen for maintaining laboratory beetle populations.

## DATA AVAILABILITY

All raw sequencing reads used in this study are available at BioProject ID 1405530 via NCBI. Genome assemblies, annotations, scripts, and associated files are available through Figshare: https://figshare.com/projects/Hidden_in_the_genomic_bycatch_insights_into_genome_reorganization_and_population_structure_of_the_parasitic_nematode_i_Contortylenchus_reversus_i_/279578

## Notes

### Competing Interest Statement

The authors have declared no competing interest.

