## Supplemental figures and tables for "Hidden in the genomic bycatch: insights into genome reorganization and population structure of the parasitic nematode *Contortylenchus reversus*"

SUPPLEMENTARY MATERIAL

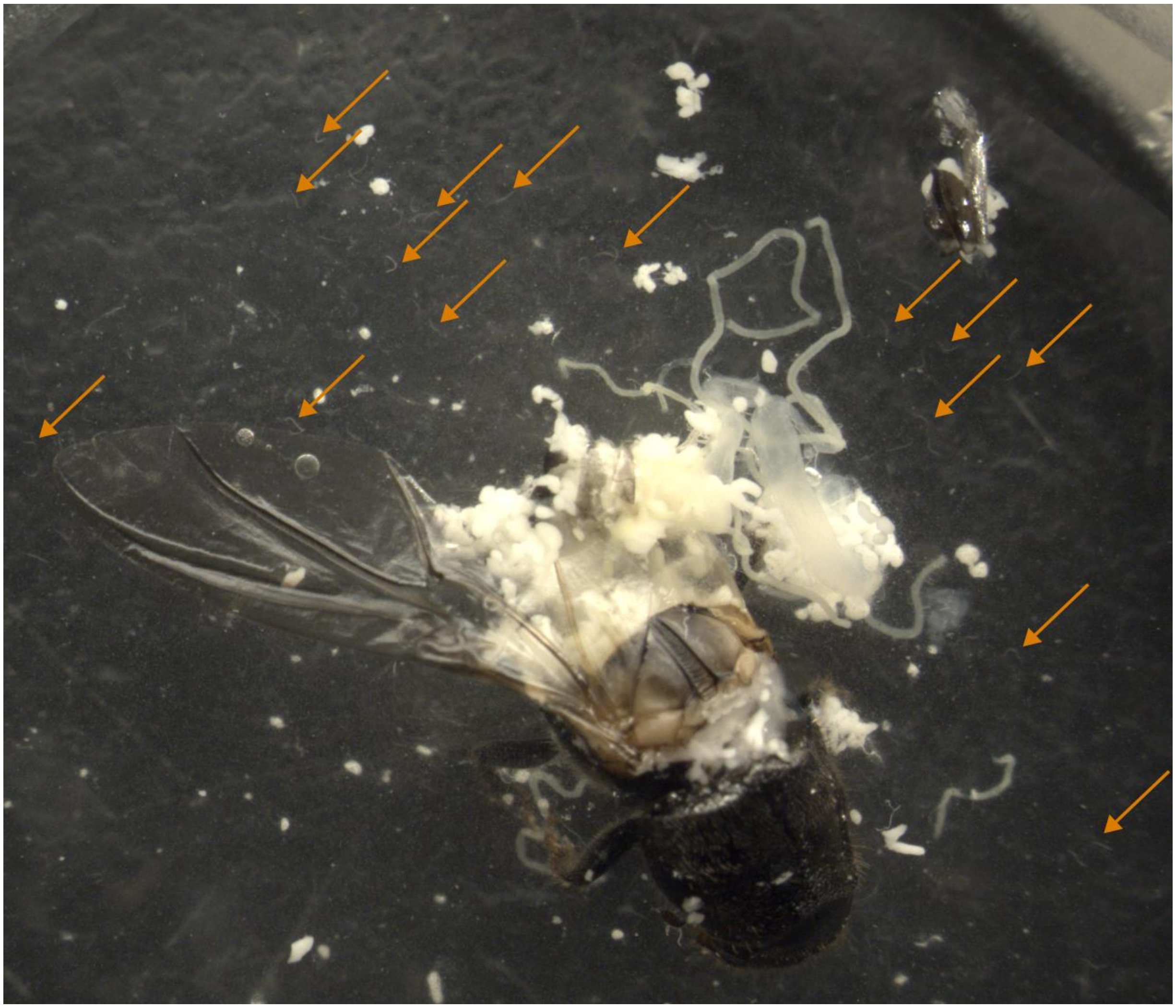

**Supplemental Figure 1. Parasitic nematode *Contortylencus reversus* in high abundance in the hemocoel of a mountain pine beetle *Dendroctonus ponderosae*.** Pictured is a dissected beetle (head oriented to bottom right of image) where the wing covers (elytra) have been removed, and the abdomen ruptured to release parasitic nematodes into a phosphate buffer solution. Orange arrows highlight several juvenile nematodes.

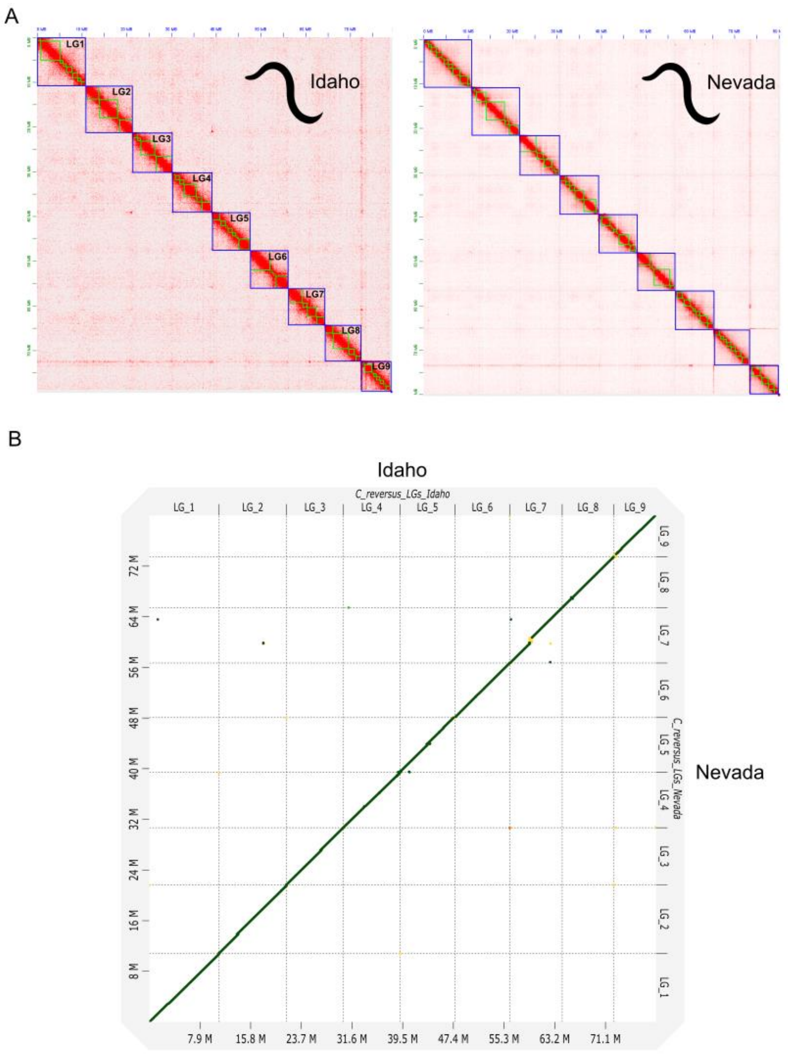


**Supplemental Figure 2. Two *Contortylenchus reversus* genomes recovered during assembly of the target host beetle, *Dendroctonus ponderosae*.** A) Hi-C contact map of *C. reversus* contigs (green boxes) recovered from the assembly and scaffolding of a male beetle from Idaho (primary reference assembly) and a female beetle from Nevada. Blue boxes outline the nine scaffolded linkage groups which appear to represent distinct chromosomes. B) Whole genome alignment between the independently assembled Idaho and Nevada assemblies shows only minor differences and overall broad collinearity.

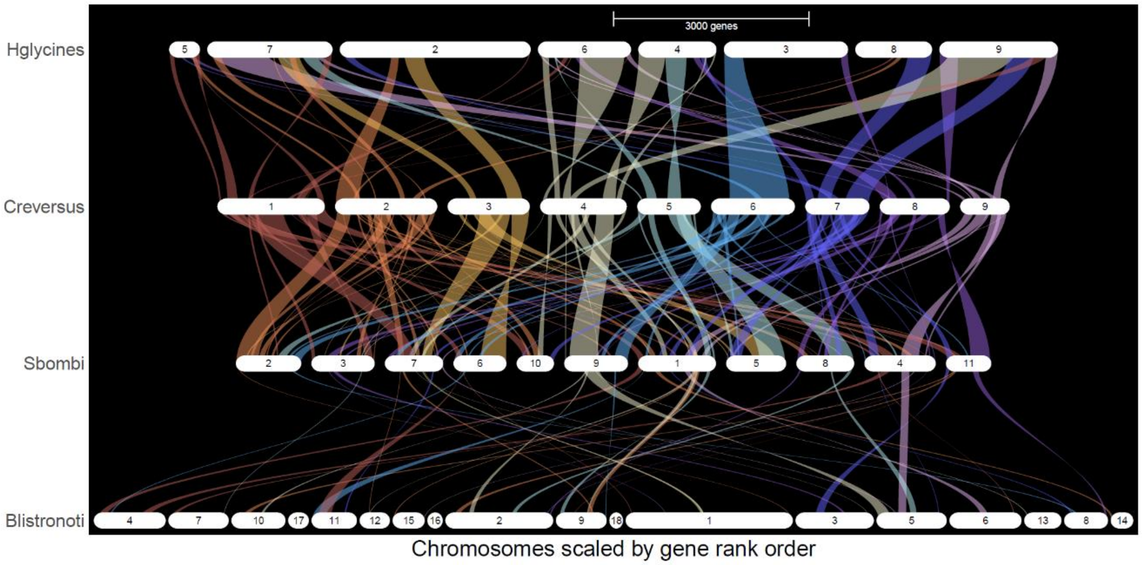


**Supplemental Figure 3. Loss of synteny among nematodes across the Tylenchoidea superfamily.** GENESPACE riparian plot of the soybean cyst nematode *Heterodera glycines*, bark beetle parasite *Contortolynchus reversus* (plotted as reference), bumblebee parasite *Sphaerularia bombi*, and the carrot weevil parasite, *Bradynema listronoti*. Note that *H. glycines*, *C. reversus*, and *S. bombi* are considered chromosome-level while *B. listronoti* is only scaffold. Ribbons connect regions of underlying genome similarity based on gene content with chromosome (or scaffold) lengths scaled by gene rank order. Extensive changes in collinearity across nematodes can contribute to the breakdown of GENESPACE synteny detection.

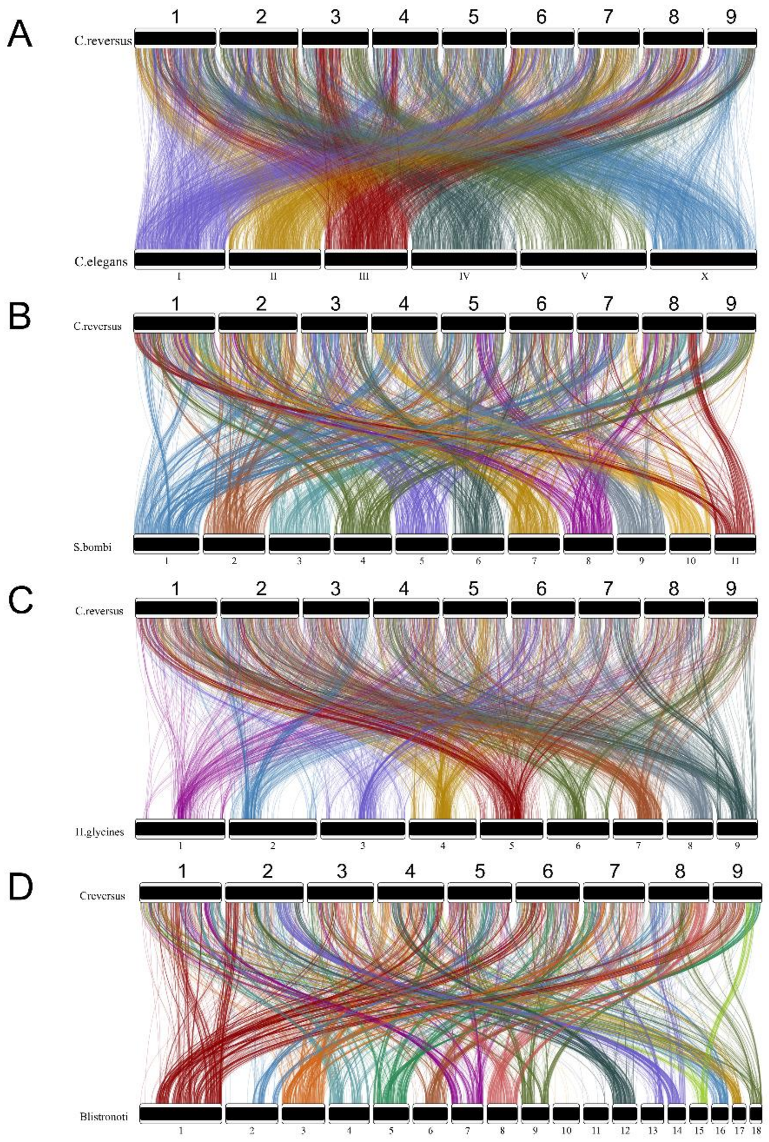


**Supplemental Figure 4**. Relative genome coordinates of 1:1 orthologs between *Contortylenchus reversus* and A) *Caenorhabditis elegans* (5,438 1:1 orthologs), B) *Sphaerularia bombi* (5,385 1:1 orthologs), C) *Heterodera glycines* (3,629 1:1 orthologs), and D) *Bradynema listronoti* (4,543 1:1 orthologs). Note that colors do not represent syntenic chromosomes or 1:1 orthologs between *C. elegans*, *H. glycines*, and *S. bombi*. The *B. listronoti* assembly is only scaffold-level. Note slight differences in overall synteny patterns can occur when comparing GENESPACE riparian plots to 1:1 ortholog plots since GENESPACE can build syntenic blocks and connections from orthogroup membership. Thus, some genes in expanded gene families can contribute to block detection in GENESPACE that would drop out of 1:1ortholog analyses. In contrast, GENESPACE requires some gene collinearity to establish syntenic blocks while plots of relative locations of 1:1 orthologs are entirely independent of neighboring gene locations in other genomes.

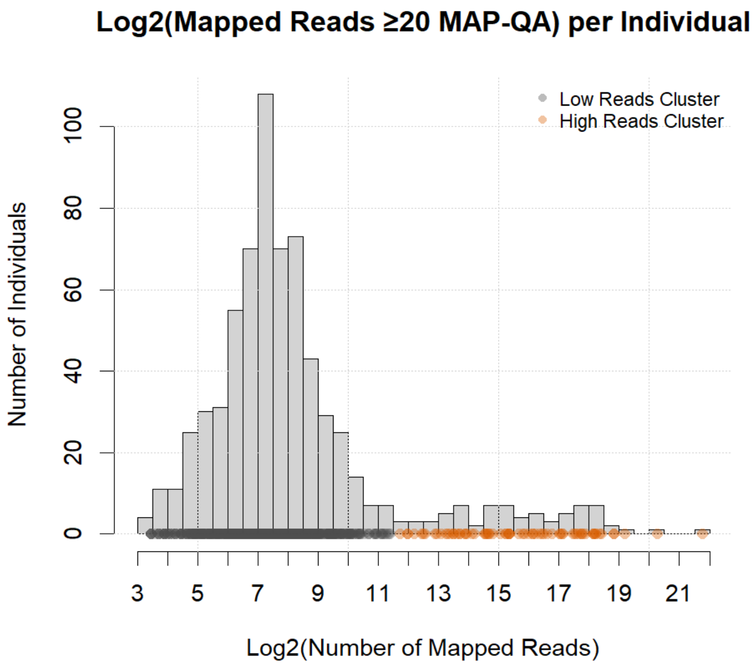


**Supplemental Figure 5**. K-means clustering to identify nematode-infected beetles. Histogram of mapped reads (log2 scale) to the *C. reversus* genome assembly for each beetle (n = 707); points along the x-axis are colored by k-means cluster assignment to two possible states, infected or uninfected (k = 2). Gray points denote the low-read cluster and orange for the high-read cluster. Based on clustering, we chose a conservative cutoff of 12 (n = 70 beetle samples with ≥ 4,096 mapped reads to the nematode genome).

**Supplemental Table 1.** BUSCO analyses of related nematode species using the nematoda_odb10 database (3,131 BUSCOs).

| | ***C. elegans*** | ***H. glycines*** | ***D. siricidicola*** | ***S. bombi*** | ***B. listronoti*** | ***C. reversus*** |
|---|---|---|---|---|---|---|
| **Complete** | 99.9% | 59.6% | 69.1% | 71.9% | 64.0% | 72.5% |
| **Single copy** | 99.5% | 51.5% | 67.5% | 70.4% | 50.9% | 70.8% |
| **Duplicated** | 0.4% | 8.1% | 1.6% | 1.5% | 13.1% | 1.7% |
| **Fragmented** | 0.1% | 2.9% | 4.4% | 3.0% | 3.2% | 3.2% |
| **Missing** | 0.0% | 37.5% | 26.5% | 25.1% | 32.8% | 24.3% |

**Supplemental Table 2.** Genome size and gene density comparison of *C. reversus* with other related nematode species.

| **Species** | **Genome size (Mb)** | **Total protein-coding genes** | **Gene density (genes/Mb)** | **Reference** |
|---|---|---|---|---|
| *C. reversus* | 79.4 | 11,244 | 141.6 | This study |
| *C. elegans* | 100.3 | 19,984 | 199.3 | GCA_000002985.3 |
| *S. bombi* | 109.1 | 11,673 | 107.0 | GCA_964235305.1 |
| *H. glycines* | 158.0 | 22,465 | 142.3 | GCA_004148225.2 |
| *B. listronoti* | 80.6 | 17,886 | 221.8 | GCA_024678965.1 |

**Supplemental Table 3.** Number of *Dendroctonus ponderosae* beetles per collection location determined to be infected with *C. reversus*. Location naming scheme as in Dowle et al. (2017).

| **Location** | **Infected** | **Not infected** | **Total** |
|---|---|---|---|
| AZ1 | 10 | 10 | 20 |
| AZ2 | 5 | 15 | 20 |
| CA1 | 0 | 20 | 20 |
| CA2 | 0 | 18 | 18 |
| CA3-4 | 10 | 22 | 32 |
| CA5 | 0 | 20 | 20 |
| CA6 | 0 | 20 | 20 |
| CAN1 | 0 | 20 | 20 |
| CAN2 | 0 | 20 | 20 |
| CO | 1 | 19 | 20 |
| ID1 | 0 | 20 | 20 |
| ID2 | 0 | 20 | 20 |
| ID3 | 3 | 16 | 19 |
| ID4 | 7 | 13 | 20 |
| ID5 | 4 | 16 | 20 |
| MT | 0 | 16 | 16 |
| MX | 2 | 16 | 18 |
| NV1 | 3 | 17 | 20 |
| NV2 | 1 | 18 | 19 |
| NV3 | 2 | 18 | 20 |
| NV4 | 0 | 20 | 20 |
| NV5 | 5 | 15 | 20 |
| NV6 | 0 | 20 | 20 |
| NV7 | 0 | 20 | 20 |
| NV8 | 0 | 20 | 20 |
| OR1 | 0 | 18 | 18 |
| OR2 | 2 | 18 | 20 |
| OR3 | 1 | 15 | 16 |
| OR4 | 3 | 17 | 20 |
| SD | 4 | 14 | 18 |
| UT1 | 0 | 20 | 20 |
| UT2 | 0 | 19 | 19 |
| UT3 | 6 | 14 | 20 |
| WA1-2 | 0 | 35 | 35 |
| WY | 1 | 18 | 19 |
| **Sum** | **70** | **637** | **707** |

**Supplemental Table 4.** Nucleotide diversity (π) of *Contortylenchus reversus* using all-sites.

| Chromosome | Average π | Number of Sites |
|---|---|---|
| **LG_1** | 0.00034085 | 66537 |
| **LG_2** | 0.00041825 | 69695 |
| **LG_3** | 0.00025369 | 49402 |
| **LG_4** | 0.00041237 | 54691 |
| **LG_5** | 0.00038739 | 50589 |
| **LG_6** | 0.00031425 | 47474 |
| **LG_7** | 0.00029183 | 48089 |
| **LG_8** | 0.0002689 | 47056 |
| **LG_9** | 0.00025754 | 37191 |

**Supplemental Table 5.** Nucleotide diversity (π) of *D. ponderosae* using all-sites.

| Chromosome | Average π | Number of Sites |
|---|---|---|
| **Chr1** | 0.00345151 | 343627 |
| **Chr2** | 0.00498882 | 312439 |
| **Chr3** | 0.00720487 | 239565 |
| **Chr4** | 0.00512998 | 262866 |
| **Chr5** | 0.00485859 | 262112 |
| **Chr6** | 0.00436164 | 265850 |
| **Chr7** | 0.00455612 | 256453 |
| **Chr8** | 0.00496669 | 194525 |
| **Chr9** | 0.0051977 | 160211 |
| **Chr10** | 0.00551704 | 154448 |
| **Chr11** | 0.00572233 | 51562 |
